# Nonlinear trajectories of language network development

**DOI:** 10.64898/2026.03.25.714106

**Authors:** Wenjing Yu, Ping Ju, Xiaohong Yang, Elizabeth Jefferies, Xi-Nian Zuo

**Affiliations:** Department of Psychology, Renmin University of China, No.59, Zhongguancun Street, Beijing, 100872, China; Jiangsu Collaborative Innovation Center for Language Ability, Jiangsu Normal University, No. 101, Shanghai Road, Xuzhou, 221100, China; Department of Psychology, University of York, Heslington, York, YO105DD, UK; State Key Laboratory of Cognitive Neuroscience and Learning, Beijing Normal University, No. 19, Xinjiekouwai Avenue, Beijing, 100875, China; Developmental Population Neuroscience Research Center, IDG/McGovern Institute for Brain Research, Beijing Normal University, No. 19, Xinjiekouwai Avenue, Beijing, 100875, China; National Basic Science Data Center, No. 2, Dongsheng South Road, Beijing, 100190, China

## Abstract

How the human brain organizes complex cognitive functions remains unresolved, particularly regarding the debate between localized and distributed architectures. Here, we show that the language network undergoes a non-linear developmental reorganization that reconciles these views. Using multimodal neuroimaging and behavioral measures, we identify a three-stage trajectory: early localization, a transiently distributed state during adolescence marked by a connectivity dip, and a return to refined localization in adulthood. This adolescent dip is behaviorally meaningful and contributes to integrative network architecture. Convergent shifts in functional connectivity and brain–behavior relationships identify adolescence as a critical window for large-scale network remodeling. Our findings provide a unifying framework for language network development and suggest that transient redistribution may represent a general principle of human brain maturation.

## Main Text

A fundamental question in developmental cognitive neuroscience is how complex cognitive functions, such as language comprehension and reading, become organized in the brain over the course of development. From childhood to adulthood, the brain undergoes profound structural and functional changes, yet the underlying principles governing this dynamic reorganization remain a subject of intense debate. The language system, supported by distributed yet functionally constrained cortical systems, provides a tractable case for understanding how developmental change balances specialization and integration. To explain how brain networks are organized across development, two opposing theoretical perspectives have been proposed. The first perspective, referred to here as the *Developmental Localization Account*, emphasizes that cognitive processing in childhood relies on relatively diffuse and broadly distributed networks (1, 2). With increasing age, this diffuse pattern gradually contracts and reorganizes into networks that are more spatially constrained and functionally specialized, reflecting a shift from broadly distributed to localized systems that support efficient information processing (3–5). This theory anticipates that children typically exhibit diffuse and less differentiated functional connectivity (FC), with development marked by increasing segregation and selective strengthening of connections within networks as they become more specialized and efficient. In contrast, a second perspective, referred to below as the *Developmental Distribution Account*, argues that FC in childhood is initially supported by relatively localized systems, which gradually develop into more distributed cross-system connection patterns during adolescence and adulthood (6–9). According to this view, the defining characteristic of brain maturation lies in the enhancement of cross-network integration associated with the flexible allocation of cognitive resources (7, 10, 11). From this perspective, this account anticipates that children show relatively constrained cross-network interactions, whereas adolescence and adulthood are characterized by stronger integration across functional systems.

A central limitation of these traditional developmental models is their reliance on linear assumptions, which struggle to capture the non-monotonic nature of brain maturation. While accounts emphasizing specialization or integration have both advanced our understanding of brain organization, they typically presume a continuous trajectory toward a single mature configuration. Converging evidence instead points to adolescence as a distinct phase of heightened plasticity and large-scale reorganization, marked by transient reductions in network efficiency, synaptic pruning, and widespread changes in functional architecture, which have been proposed to support behavioral variability and cognitive exploration (5, 10, 12–14). Consistent with this view, Liang et al. (2025) reported that whole-brain network variability peaks during adolescence, average functional connectivity reaches a minimum in early adolescence, and modularity and small-world properties continue to mature into adulthood.

### Scope of The Study

Building on these insights, we propose a unified, three-stage developmental model that reconciles these seemingly contradictory localization and distribution accounts. We hypothesize that the functional organization of specialized cognitive networks follows a “**localization-to-distributed reorganization-to-refined localization**” trajectory: (1) Childhood is characterized by a relatively localized organization, where processing relies on coarse, short-range connections within nascent, functionally specific modules. (2) Adolescence serves as a pivotal reorganization phase, transitioning into a distributed organization. This stage is marked by a strategic, albeit temporary, expansion of network scope and a potential weakening of some local connections to facilitate cross-system integration, cognitive flexibility, and the exploration of new computational strategies. (3) Adulthood involves refined localization, where networks reconfigured into a highly efficient, specialized, and streamlined architecture, with selectively strengthened task-critical pathways and pruned redundant ones (Fig. 1A).

**Fig. 1.**
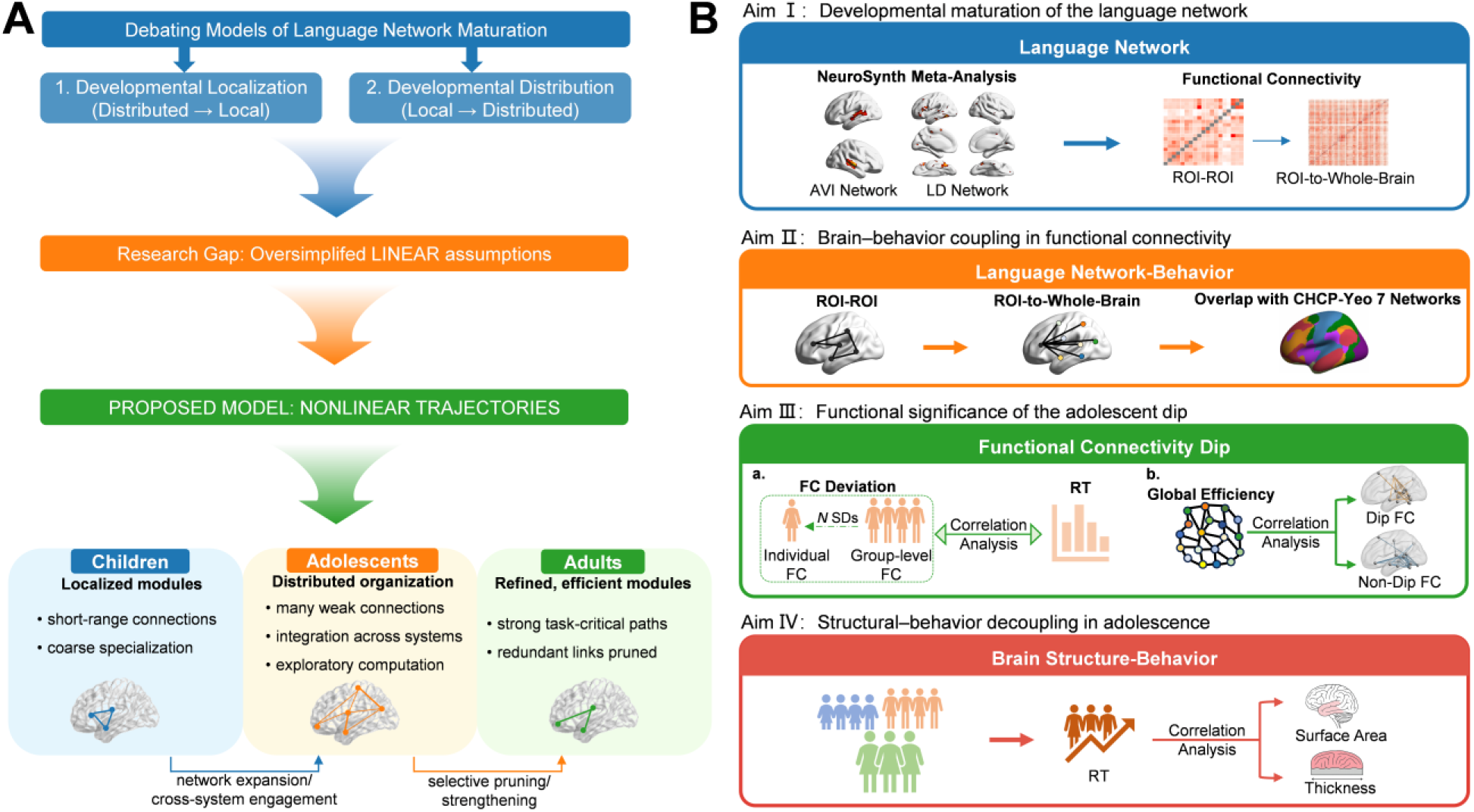
Theoretical model and data-analysis workflow. Panel **A** illustrates the theoretical model proposed in this study. From childhood to adolescence and into adulthood, the functional organization of specialized cognitive networks is proposed to follow a “localization-distributed reorganization-refined localization” trajectory. The model is tested using two tasks: audiovisual integration (AVI) and lexical decision (LD). Panel **B** summarizes the data-analysis workflow. We first conducted ROI-to-ROI, ROI-to-whole-brain analyses within the language network. Next, we examined how developmental changes in the language network relate to behavioral performance using both ROI-to-ROI and ROI-to-whole-brain approaches, and evaluated the overlap between significant functional connections and large-scale brain networks. We then focused on the adolescent reorganization phase, specifically investigating the relationship between the adolescent connectivity dip and behavioral performance, as well as the cognitive significance of this dip. Finally, we assessed associations between structural features (cortical thickness and surface area) and behavioral measures.

To test the proposed three-stage developmental model, we collected a language-focused developmental cohort spanning from childhood to adulthood as part of the Chinese Color Nest Project (CCNP). This cohort includes language-related tasks alongside resting-state and structural neuroimaging measures, enabling a systematic test of the model. Detailed information about CCNP is provided in the Participants section of the Supplementary Materials. In addition, we designed a multi-level analytical framework that moves beyond single metrics to capture the dynamic interplay between neural architecture, cognitive behavior, and structural constraints (Fig. 1B). First, to track the intrinsic maturation of the language network, we meta-analytically defined two sets of language-relevant regions-of-interest (ROIs) a priori in NeuroSynth: one associated with audiovisual integration processes that support our capacity to link orthographic and phonological information (NeuroSynth meta-analysis for the term “audiovisual”), and a second associated with lexical–phonological processing involved in semantic access (NeuroSynth meta-analysis for the term “lexical decision”). We then quantified developmental trajectories through ROI-to-ROI resting-state FC and ROI-to-whole-brain FC, predicting that the connectivity strength within these language networks would progressively increase as they mature. During adolescence, however, FC strength may show greater variability, including temporary decreases or developmental plateaus, reflecting ongoing network reorganization.

Second, to directly link these neural changes to cognitive maturation, we correlated FC strength with behavioral performance across development. Specifically, behavioral performance was indexed by RT difference, with smaller values indicating more efficient processing. We predicted a three-stage shift in the spatial pattern and functional direction of brain-behavior coupling: from sparse, localized, and efficient (negative correlations) in childhood, to widespread but inefficient coupling in adolescence, a stage in which stronger connectivity may paradoxically predict worse performance (positive correlations), reflecting a transitional state of network reconfiguration in which broader connectivity recruitment does not yet translate into efficient processing. In adulthood, we anticipated a return to a refined, localized, and efficient pattern (negative correlations). To understand the system-level manifestation of network expansion and refinement, we also projected significant functional connections onto large-scale brain networks. We predicted a progressive shift toward greater involvement of control- and attention-related systems, specifically the frontoparietal, dorsal attention, and ventral attention networks, from childhood to adulthood, consistent with increasing efficiency in cognitive control.

Third, to move beyond group-level averages, we examined whether the adolescent reorganization has meaningful functional consequences using two complementary analyses. First, we asked whether individual differences in the magnitude of the adolescent connectivity dip (i.e., the transient reduction in FC strength) are related to behavioral performance, using a developmentally anchored deviation analysis. Second, we assessed the system-level consequences of this reorganization by testing how connections that show a dip, versus those that do not, contribute to global network efficiency during adolescence. Together, these analyses determine whether the connectivity dip reflects nonspecific inefficiency or a structured reconfiguration that supports the emergence of mature network integration.

Finally, to test the hypothesis that adolescence is marked by a structural decoupling from behavior, a transition that may permit the large-scale functional reorganization observed during this period, we investigated the relationship between structural properties (cortical thickness, surface area) and behavioral performance. We predicted strong structure-behavior coupling in childhood, reflecting initial reliance on structural scaffolding; attenuated coupling in adolescence, coinciding with functional network redistribution; and a lack of such coupling in adulthood, indicating a transition to a functionally optimized state.

By dissecting the language network into its core subsystems and applying this multi-level analytic approach, we transcend the oversimplified “localization vs. distribution” debate. We test the overarching hypothesis that the language network undergoes a three-stage nonlinear developmental trajectory, which may reflect a fundamental principle of neurocognitive optimization.

## Results

### Behavioral strategies shift from perceptual processing to cognitive control during development

We first examined developmental changes in behavioral performance during audiovisual integration and lexical decision tasks. RTs decreased significantly with age in both tasks across all conditions (all ps < .001), indicating progressive gains in processing efficiency (see Fig. S1). The RT difference (incongruent–congruent for audiovisual integration; pseudoword–real word for lexical decision) exhibited distinct patterns across tasks. In the audiovisual integration task, the RT difference decreased linearly with age (edf = 1.00, F = 4.32, p < .05), reflecting a reduction in the cost of processing cross-modal speech conflict. In contrast, the lexical decision task showed no significant age-related change in RT difference (edf = 1.00, F = 0.15, p = .70), suggesting a more stable developmental profile for lexical-semantic discrimination.

More importantly, the strategic basis of performance underwent a fundamental shift: in childhood, RT differences between conditions were correlated with both easy and hard conditions, suggesting undifferentiated perceptual processing. By adolescence and adulthood, however, performance differences were driven exclusively by the more demanding conditions (see Fig. S2), signaling the emergence of selective, control-dependent processing strategies. Task-specific results are reported in Section 2.1 of the Supplementary Text.

### Core language connectivity reveals trajectories instantiating three-stage development

ROI–ROI analyses revealed heterogeneous developmental modes (Fig. 2A-C). We first examined the connections unique to each network (i.e., excluding the shared hubs analyzed separately in Fig. 2C). The audiovisual integration network (Fig. 2A) was shaped by linear increases (57.14%) and adolescent dips (35.72%). The lexical decision network (Fig. 2B) was dominated by linear increases (81.58%), with smaller proportions showing adolescent dip (7.89%), developmental plateau (7.89%), or linear decrease (2.64%). Beyond the developmental patterns within each network, we sought to characterize the maturation of the functional integration between the two language subsystems. We specifically examined the FC between the audiovisual integration and lexical decision networks (Fig. 2C). The developmental trajectories of these cross-network connections were striking: while linear increases remained the most common pattern (49.02%), the proportion of connections exhibiting an “adolescent dip” was markedly higher (43.14%) than that observed within either network individually. This pattern highlights that dynamic reorganization during adolescence is particularly prominent in the integrative circuitry linking distinct language subsystems. For all sub-analyses, a consistent developmental gradient in FC strength was observed, with adults demonstrating the strongest connectivity, followed by adolescents and then children.

**Fig. 2.**
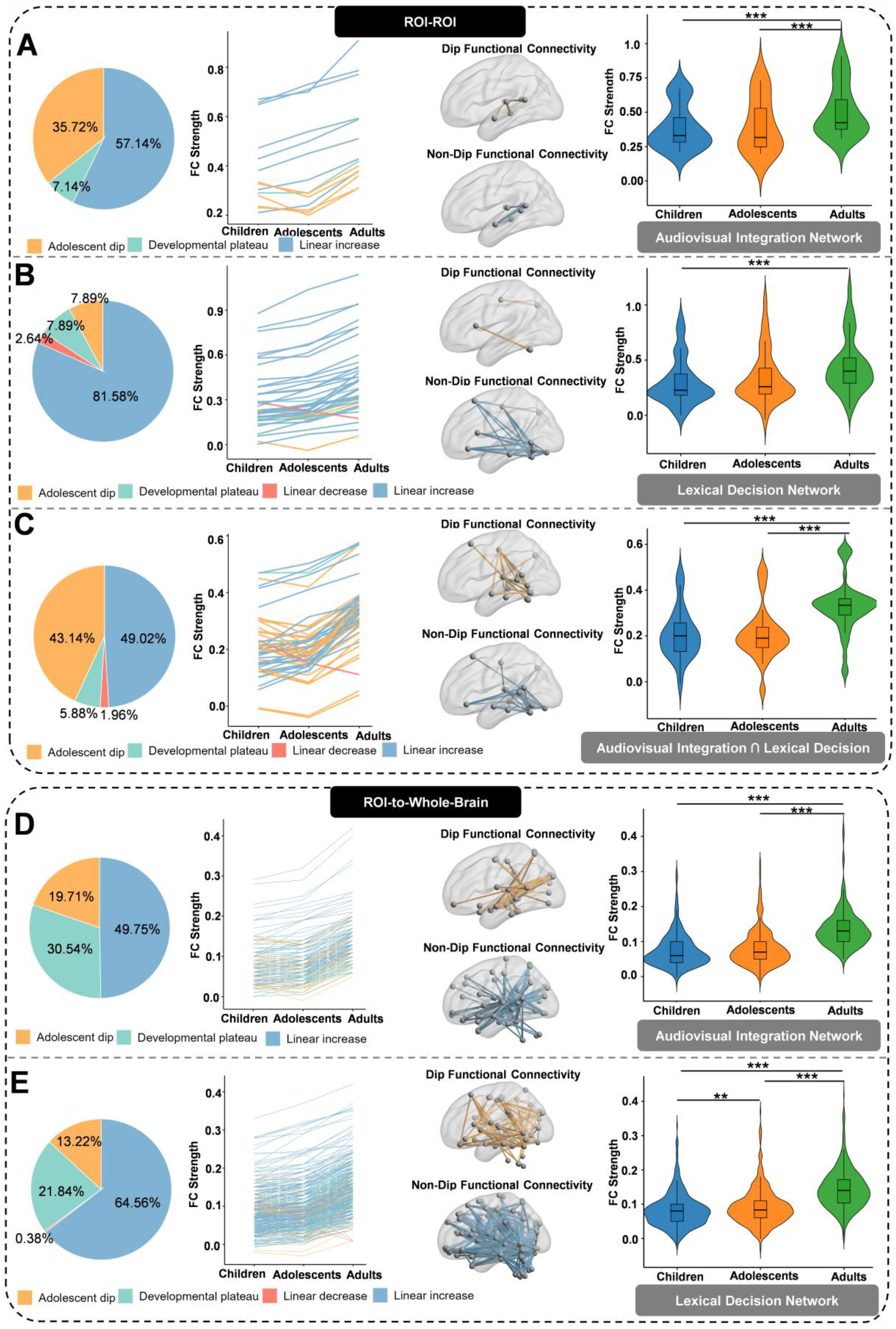
FC results of the language network. Panels A–C show the ROI–ROI FC patterns within the audiovisual integration network, within the lexical decision network, and between the two networks, respectively. Panels D and E display the whole-brain FC distributions using the audiovisual integration and lexical decision networks as seeds, respectively. The pie charts depict the proportions of different developmental trajectories, and the line graphs show the temporal patterns of each trajectory; both share a common legend. The brain maps visualize the spatial distribution of connections showing significant developmental changes, with orange and blue indicating adolescent dip and non-dip trajectories (i.e., developmental plateau, linear decrease, and linear increase), respectively. Finally, the violin plots indicate group differences in FC strength. Note: **p < .01; ***p < .001.

Seed-based whole-brain analyses confirmed that the developmental trajectories observed within the core language network reflect a broader principle of brain maturation. Across both seeds, linear increases constituted the dominant pattern, accompanied by a secondary feature of a transient decline and plateau during adolescence (Fig. 2D and 2E). Adults exhibited the strongest whole-brain connectivity for both seeds (ps < .001).

In summary, the language network matures through a dynamic process of progressive strengthening of essential pathways and selective refinement of others during adolescence, ultimately crystallizing into the highly efficient, specialized architecture observed in adulthood.

### Behaviorally-relevant connectivity undergoes a three-stage reorganization

In childhood, brain-behavior correlations were few and restricted. In the lexical decision task, ROI-ROI analysis revealed that stronger connectivity between the left ACC and left inferior frontal regions (IFG orbital and opercularis) was associated with smaller RT differences (i.e., higher processing efficiency) (r = −0.30, p = .023; r = −0.29, p = .026; Fig. 3A). This localized, efficient pattern was complemented by ROI-whole-brain findings (Fig. 3B): in the audiovisual integration task, a single connection [left TPOJ1–left superior parietal lobule (SPL)] was negatively correlated with the RT difference (r = −0.52, p < .001), as was a connection in the lexical decision task [left MTG–left middle frontal gyrus (MFG); r = −0.62, p < .001].

In adolescence, the number of behaviorally-relevant connections surged. However, this expansion was inefficient. In the lexical decision task, ROI-ROI analysis showed that stronger ACC–SMA connectivity was linked to larger RT differences (i.e., lower efficiency) (r = 0.23, p = .023; Fig. 3A). This pattern was robustly confirmed at the whole-brain level (Fig. 3B), where all significant connections in both tasks showed positive correlations with RT difference [e.g., right A1–left middle occipital gyrus (MOG), r = 0.37, p < .001], meaning stronger connectivity was associated with a greater cost in resolving cross-modal conflict or lexical uncertainty.

In adulthood, the language network reconfigured into a streamlined and efficient state, with a selective re-localization of function. In the audiovisual integration task, ROI-ROI analysis identified the left TPOJ1 as a mature hub (Fig. 3A), with its stronger connectivity to bilateral STV and left STSda supporting more efficient integration (reflected by smaller RT differences) (r = −0.36 to −0.30, ps < .06). Similarly, in the lexical decision task (Fig. 3A), stronger connectivity between temporal regions (right ITG–left PHG) was associated with a reduced pseudoword cost (r = −0.24, p = .049). This return of efficient negative correlations was also evident at the whole-brain level [e.g., right STV–left superior frontal gyrus (SFG), r = −0.31, p = .007 for the audio-visual task; left MTG–left ITG, r = −0.64, p < .001 and left ITG–right ITG, r = −0.44, p < .001 for the lexical decision task; Fig. 3B].

This three-stage pattern was corroborated at the system level (Fig. 4B and 4C). Mapping significant connections onto the CHCP-derived Yeo seven-network atlas revealed that in both tasks, the control and attention networks showed progressively increasing engagement with age. However, the specific attentional subsystems driving this engagement were task-dependent: ventral attention networks dominated audiovisual integration (adults: 11%), while dorsal attention networks specialized for lexical decision (adults: 46%). Notably, adolescence was characterized by exuberant network recruitment, with total voxel counts (audiovisual integration: n = 16186; lexical decision: n = 21290) significantly exceeding that in the adult (n = 3868; n = 3720) and child groups (n = 3444; n = 1603). This hyper-engagement suggests adolescence acts as a period of expansive network exploration preceding adult refinement. Detailed statistical results are provided in Section 2.2 of the Supplementary Text.

**Fig. 4.**
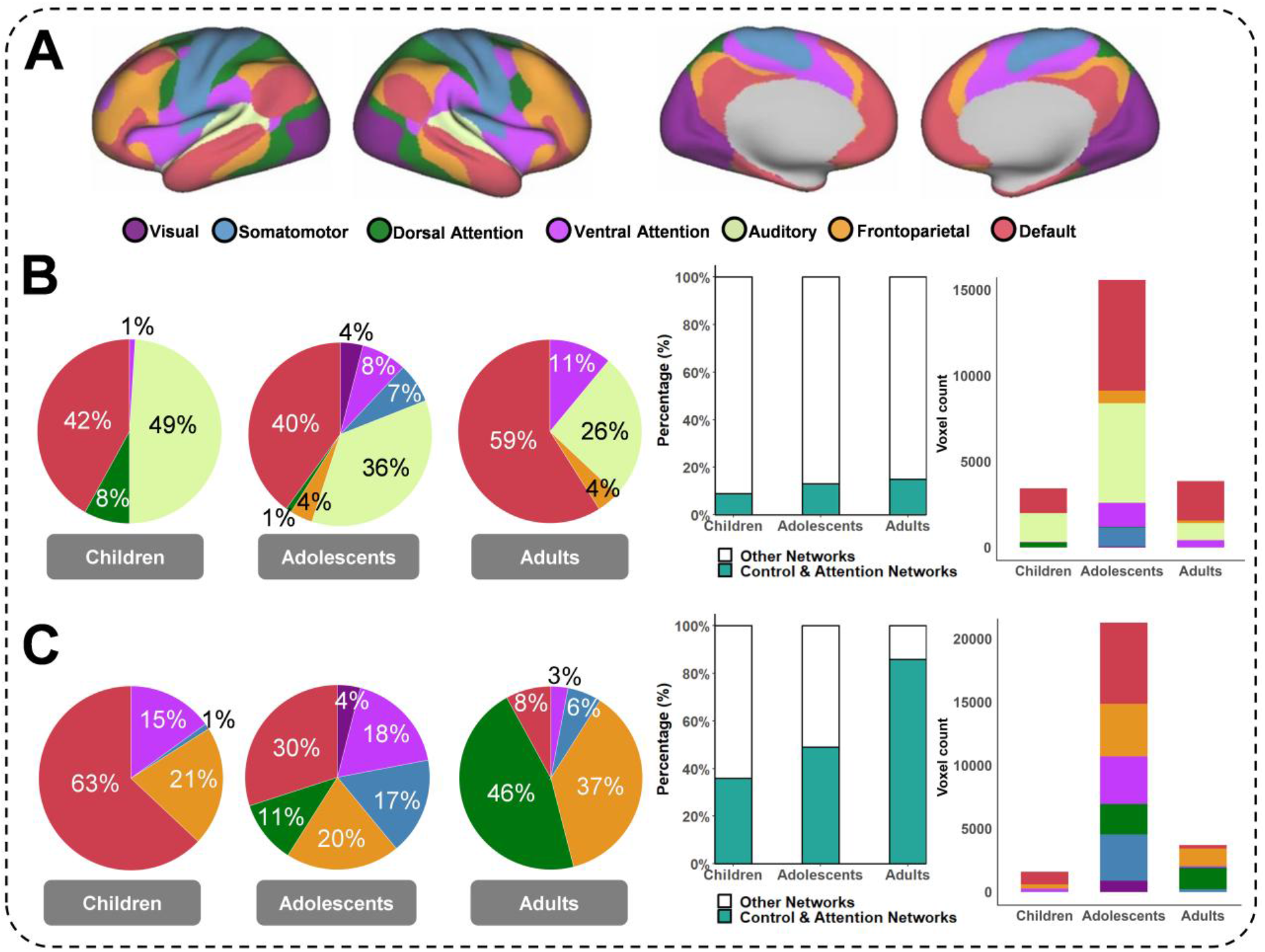
Results of large-scale brain networks. Panel A presents a schematic illustration of the seven large-scale brain networks based on the CHCP-derived Yeo 7-network parcellation. Panels B and C respectively depict the overlap between significantly connected regions and these large-scale brain networks in the audiovisual integration task and the lexical decision task. From left to right, the pie charts illustrate the distribution of large-scale networks in children, adolescents, and adults. The stacked bar chart on the left summarizes the percentage of age-related changes in the involvement of the control and attention networks (i.e., the dorsal attention, ventral attention, and frontoparietal networks), while the stacked bar chart on the far right quantifies the voxel counts within each large-scale network for each age group.

### Successful implementation of the adolescent dip relates to behavioral advantage and reduced transient over-distribution

To further elucidate the functional significance of the adolescent dip observed at the group level, we conducted two complementary analyses to clarify how the dip relates to immediate task performance and why it might be a necessary feature for the development of an optimized, adult-like network architecture.

Individual deviation analysis revealed task specific associations between FC deviation scores and performance (Fig. S3A and S3B). In the audiovisual integration task (Fig. S3A), a significant positive correlation emerged in adolescents (r = 0.201, p = .041), where greater positive deviation from the adolescent mean (i.e., higher-than-average FC, indicating a less pronounced dip) was associated with larger RT differences and reduced behavioral efficiency, whereas greater negative deviation (i.e., lower-than-average FC, reflecting a stronger dip) correlated with smaller RT differences and enhanced efficiency. In adults, this reversed to a significant negative association (r = −0.261, p = .022): greater negative deviation impaired performance, highlighting the emergence of a stabilized, efficient architecture. Children exhibited no association (r = −0.050, p = .673). In contrast, the lexical decision task showed a similar directional pattern but no significant correlations across groups (all ps > .27), consistent with its predominantly linear and stable maturation for semantic discrimination (Fig. S3B). These findings highlight the adolescent dip as a critical vulnerability window, particularly in perceptual integration subsystems, where deviations toward less “dipped” states (positive offsets) incur substantial behavioral costs.

Dip contribution analysis further clarified the network level role of dip connections (Fig. S3C and S3D). In adolescents, both Dip and Non-dip FC showed strong positive correlations with global network efficiency (AV task: Dip r = 0.336, p < .001; Non-dip r = 0.651, p < .001. LD task: Dip r = 0.597, p < .001; Non-dip r = 0.730, p < .001). This indicates that both sets of connections ultimately support the formation of the adult-like integrative architecture, demonstrating that adolescent dips do not indicate functional degradation. In addition, dip connections contribute more modestly to efficiency, consistent with their role as reorganization-sensitive pathways that transiently weaken during adolescence to permit large-scale redistribution before being reincorporated into the mature network.

Together, these findings show that the adolescent dip functions both as a group-level marker of distributed reorganization and as an individual-level determinant of behavioral efficiency. Individuals who align more closely with the adolescent-specific remodeling state demonstrate better task performance, and both dip and non-dip connections contribute to the integrative network architecture that supports mature language processing. These results reinforce the view that adolescent redistribution is a necessary precursor to the emergence of refined, adult-like integration.

### Structural decoupling parallels functional reorganization

Finally, we tested the predicted developmental shift in structure–behavior coupling. In the audiovisual integration task, children exhibited significant negative correlations between RT difference and cortical surface area in two core regions: right STSvp (r = −0.34, p = .004) and right STV (r = −0.34, p = .003); larger surface area was associated with smaller (more efficient) behavioral costs (Fig. S4A). In the lexical decision task, children showed a marginal positive correlation in left MTG (r = 0.25, p = .053), suggesting that reduced surface area was tentatively linked to lower pseudoword costs (Fig. S4B). These structure–behavior associations largely disappeared in subsequent stages. In adolescence, the only surviving correlation was a negative relationship in the lexical decision task between left fusiform gyrus surface area and RT difference (r = −0.26, p = .011; Fig. S4C); no correlations were observed in the audiovisual integration task. In adulthood, structure–behavior coupling was entirely absent in both tasks (all ps > .20).

Thus, robust structure–behavior relationships present in childhood progressively weaken and ultimately vanish across development. These findings are consistent with a developmental transition in which early reliance on structural properties gives way to greater functional autonomy. Such decoupling may permit the large-scale functional reconfiguration observed during adolescence, thereby facilitating the eventual emergence of the efficient, specialized language architecture seen in adulthood. No significant correlations were observed between cortical thickness and behavioral measures in any region across all three age groups and both tasks after correction for multiple comparisons.

## Discussion

This study systematically characterizes the developmental trajectory of language network organization from childhood to early adulthood across behavioral, functional, and structural levels. Using data from the CCNP, from which we derived a language-focused cohort, we demonstrate that the language network follows a nonlinear three-stage developmental pattern: from locally scaffolded processing in childhood, to a distributed, reconfigured, and transiently costly state in adolescence, and finally to refined, efficient localization in adulthood, which we synthesize in the model in Fig. 5. This integrative framework not only reveals the multi-trajectory nature of language development but also highlights adolescence as a pivotal period for neural and cognitive reorganization.

**Fig. 5.**
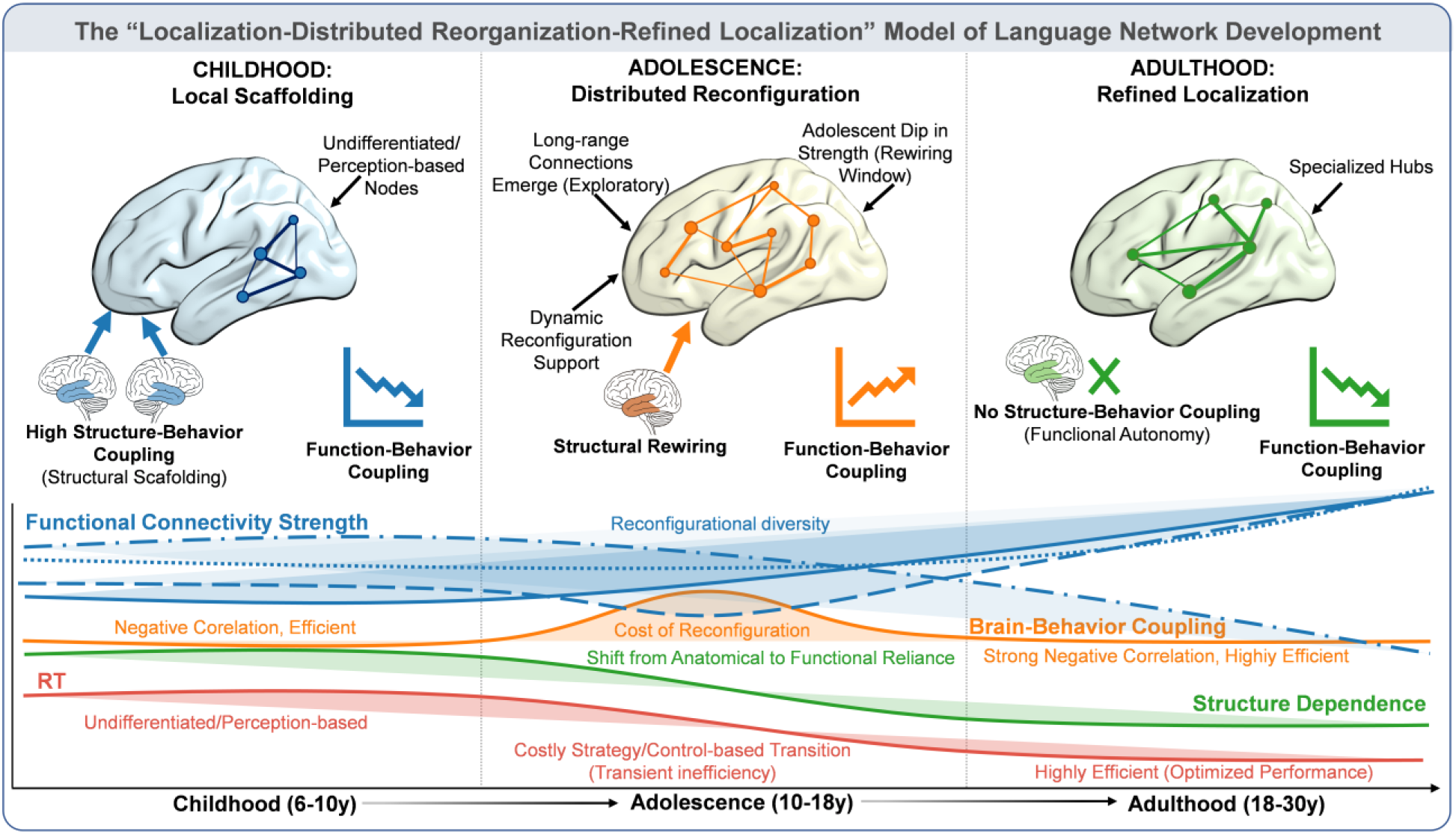
The synthesized model of language network development.

At the FC level, the study identifies adolescence as a critical period of language network reconfiguration (Fig. 5). Although overall connectivity increases with age, the connections within language subsystems, between subsystems, and between the language network and the whole brain all exhibit pronounced dips or developmental plateaus during adolescence. This pattern aligns with the surge of neural plasticity associated with synaptic pruning and functional reorganization in adolescence (10, 12, 15). These connectivity perturbations thus reflect exploratory topological remodeling rather than degeneration, supporting the cross-system shift from perceptual to control-oriented processing. This finding deepens our understanding of adolescence as a phase of network resculpting.

The three-stage pattern of function–behavior coupling further strengthens the three-stage model, as illustrated in Fig. 5. In childhood, coupling is restricted to a small set of local regions and is characterized by negative associations (i.e., stronger connectivity predicts better performance), reflecting reliance on discrete functional units (16, 17). During adolescence, coupling becomes widespread yet unstable, characterized by a shift toward positive correlations (i.e., stronger connectivity paradoxically predicts worse performance), indicative of high plasticity coupled with low processing efficiency (10, 15). In adulthood, coupling reconverges into a more efficient, localized pattern, restoring negative correlations (18). This trajectory, from localization, to more functional distribution, to refined localization, accompanied by a transitional reversal in coupling direction during adolescence, is not consistent with unidirectional developmental models that propose increasing localization or functional distribution (1, 2, 7, 10, 11) and offers a more realistic, nonmonotonic framework of brain maturation. Large-scale network analyses further show that engagement of the control and attention networks increases from childhood to adulthood, indicating that higher-order control systems play a driving role in the development of language processing (19, 20). This pattern is consistent with evidence that regions commonly associated with the control and attention networks show increased involvement in language processing after adolescence (13, 21). Thus, the maturation of the language network depends not only on internal reorganization but also on cross-system coordination with executive-control networks.

The individual deviation and dip-contribution analyses further illuminate the functional meaning of the adolescent reorganization observed at the group level. Notably, individuals whose FC deviated toward a more “dipped” profile (i.e., lower-than-average FC in adolescence) exhibited better behavioral efficiency in the audiovisual integration task, whereas those with less dipped or elevated connectivity showed reduced performance. This suggests that the adolescent dip is not merely a sign of inefficiency, but rather a behaviorally relevant marker of network transition, where alignment with the developmental “reorganization window” predicts more adaptive processing. Moreover, the dip-contribution analysis revealed that both “dip” and “non-dip” connections contributed positively to global network efficiency during adolescence, indicating that even transiently weakened pathways participate in system-wide integration. Together, these findings refine our understanding of the adolescent dip: it represents a purposeful, network-wide reconfiguration that facilitates the shift from childhood’s locally scaffolded processing to adulthood’s refined integration, rather than a simple disruption or regression. This highlights adolescence as a critical period of functional optimization, in which network restructuring supports the emergence of mature, efficient language processing.

The structure–behavior coupling results reveal a progressive developmental decoupling (Fig. 5), complementing the functional reorganization described above. In childhood, language processing strongly depends on structural substrates such as cortical surface area; this dependence markedly decreases during adolescence and becomes restricted to only a few pathways; by adulthood, structural indices no longer predict behavioral performance. This decoupling parallels the functional reorganization observed in adolescence, suggesting that the reduction in structure– behavior coupling may permit the large-scale functional reconfiguration necessary for network redistribution (22–24).

Notably, this neurodevelopmental process is paralleled by a fundamental shift in behavioral strategies. Although the behavioral findings show that RT in both tasks decrease with age, the underlying processing strategies undergo a qualitative shift. Children rely on an undifferentiated, perception-based mode in which task difficulty effects are jointly driven by both easy and hard conditions. During adolescence, processing gradually shifts toward a control-dominated strategy primarily driven by challenging conditions, reaching stability in adulthood. This pattern suggests that the key to language development lies not in improvements in processing speed, but in a mechanistic transition from low-level perceptual dependence to goal-directed, control-based processing (3, 25, 26). Thus, the behavioral shift provides convergent evidence for the triphasic model, highlighting how cognitive strategy evolution parallels the large-scale neural reconfiguration observed across development.

Taken together, the present findings map coherently onto the triphasic nonlinear developmental model summarized in Fig. 5. First, ROI-to-ROI and seed-to-whole-brain analyses revealed a predominantly linear strengthening of FC with age, which was systematically punctuated by an adolescent dip, most notably in long-range, cross-network connections, directly substantiating the distributed reorganization phase of the model. Second, function–behavior coupling exhibited a characteristic three-stage shift: from sparse, efficient, and localized (childhood), to widespread, inefficient, and positively correlated (adolescence), and finally to reconverged, efficient, and localized (adulthood), providing a behavioral signature of the network’s sequential reorganization. Third, individual deviation analyses linked the adolescent dip directly to behavior, showing that individuals with a more pronounced dip (lower-than-average FC) exhibited better task performance in the audiovisual integration task, while graph theoretical analyses confirmed that both dip and non-dip connections contributed positively to global network efficiency during adolescence. Together, these findings indicate that the dip reflects a behaviorally relevant, purposeful recalibration of the network’s functional architecture, rather than a simple degradation. Finally, structure–behavior coupling was strong in childhood but progressively diminished across adolescence and was absent in adulthood, consistent with a developmental transition from structure-scaffolded processing to functionally optimized network integration. These multi-level results establish adolescence as a critical, reorganization-sensitive window through which the language network transitions from an early, structurally dependent architecture toward a mature, efficient, and flexibly integrated functional state.

## Conclusions

To summarize, this study proposes and validates an integrated developmental framework spanning behavioral, structural, and functional dimensions in the language network. Our findings demonstrate that language network maturation does not follow a simple monotonic trajectory, but instead unfolds through a triphasic trajectory characterized by early efficiency, adolescent network expansion, and adult functional refinement. By moving beyond linear accounts of brain maturation, we help reconcile the debate between developmental localization and distribution, demonstrating that these principles represent sequential states rather than competing endpoints. Within this trajectory, we reframe the adolescent “dip” in functional connectivity not as random or epiphenomenal developmental variation, but as an adaptive, mechanistic reconfiguration that selectively optimizes global network efficiency and individual performance. Together, these findings identify adolescence as a critical transition point where language processing shifts toward functionally autonomous, control-dependent integration. This triphasic model provides a unified theoretical perspective for understanding language development and offers a foundation for future research on large-scale network across higher-order cognitive systems.

## Acknowledgments

We thank Yang Yang for valuable discussions and insightful ideas during the early stages of the project. We also thank Zihang Zhou for her contributions to data collection, and Junhao Xu for providing support with data analysis equipment.

## Author contributions

Conceptualization: WY, XY, EJ

Methodology: WY, XY, XNZ

Cohort Investigation: XY, XNZ

Data Curation: WY, PJ

Visualization: WY, PJ, XNZ

Funding acquisition: XY, XNZ

Project administration: XY, XNZ

Supervision: XY, XNZ

Writing – original draft: WY, XY

Writing – review & editing: WY, XY, EJ, XNZ

## Data, code, and materials availability

All data supporting the findings of this study are available within the main text and/or the supplementary materials.

Additional data and code are available via OSF at https://osf.io/rcfhd/overview?view_only=86aad70db05046509ca26fbf58a3cd8c.

The cohort imaging data anonymized can be accessed via Science Data Bank as part of the Chinese Color Nest Data Community (https://ccndc.scidb.cn/en) at https://doi.org/10.57760/sciencedb.07478.

Raw data (e.g., MRI data) are available from the corresponding author upon reasonable request.

