## Supplemental Materials for "Nonlinear trajectories of language network development"

**The PDF file includes:**

Materials and Methods
Supplementary Text
Figs. S1 to S6
Tables S1 to S2

### 1 Materials and Methods

#### *1.1 Participants*

Participants were drawn from the Chinese Color Nest Project (CCNP), a lifespan brain-mind development data community of Chinese children, adolescents, and adults (*1, 2*). The present study focuses on a language-relevant subset of this dataset, including behavioral assessments and multimodal neuroimaging data (resting-state and structural scans). The sample spanned an age range from 6 to 30 years. Data were collected using a combination of accelerated longitudinal and cross-sectional designs: participants aged 6–18 years were assessed up to three times over a five-year period, with approximately 15-month intervals between sessions to minimize seasonal effects, whereas participants in early adulthood were assessed once. Behavioral assessments were administered on the day of scanning or within a few days thereafter, with consistent testing content across waves. After aligning brain and behavioral data, a total of 188 observations were obtained from 152 participants in the longitudinal sample, together with 77 observations from adult participants. Following data quality control, six participants were excluded due to excessive head motion (mean framewise displacement FD > 0.2 mm) (*3*). As a result, the final valid sample consisted of 259 participants.

According to the *Health Needs of Adolescents* report published by the World Health Organization (WHO) in 1999 and reaffirmed in 2017, adolescence is formally defined as the period between ages 10 and 19. In parallel, Piaget's theory of cognitive development suggests that the transition from the concrete operational stage (ages 7–11) to the formal operational stage (ages 12 and above) typically occurs from middle childhood to early adolescence (*4*). Therefore, in this study, participants were classified into three groups: children (aged 6.1–10 years), adolescents (10.1–18 years), and adults (18.1–30 years). For behavioral assessment, these participants completed an audiovisual integration task, a lexical decision task, or both. For each task, we further excluded participants with task accuracy below 70% or whose behavioral data were identified as outliers.

In the audiovisual integration task, after excluding four children with accuracy below 70%, as well as one child and one adult whose behavioral data were identified as outliers, valid data included 73 observations from children (*M* = 8.34 years, *SD* = 1.07; 41 males), 104 from adolescents (*M* = 12.07 years, *SD* = 1.61; 55 males), and 76 from adults (*M* = 21.70 years, *SD* = 2.29; 27 males). These participants met all quality control criteria and successfully completed the task. Participant information is shown in Fig. S5A and S5C. In the lexical decision task, a subset of participants from the audiovisual integration task took part. Specifically, 5 children, 1 adolescent, and 6 adults did not participate in the lexical decision task. After further excluding participants whose task accuracy was below 70%, the final valid sample included 59 child observations (*M* = 8.53 years, *SD* = 1.00; 35 males), 95 adolescent observations (*M* = 12.15 years, *SD* = 1.65; 48 males), and 70 adult observations (*M* = 21.70 years, *SD* = 2.23; 27 males). Participant information for this task is presented in Figs. S5B and S5C.

#### *1.2 Task paradigms*

Behavioral performance was assessed using two language tasks: an Audiovisual Integration Task and a Lexical Decision Task.

The audiovisual integration task was designed to probe the rapid and automatic mapping between visual word forms and their corresponding sounds (*5*), which forms a foundational component of reading acquisition (*6*). Participants viewed written Chinese characters while simultaneously hearing their pronunciations and indicated whether the visual and auditory inputs were matched (congruent) or mismatched (incongruent). The contrast between incongruent and congruent trials isolates the neural and cognitive costs of resolving cross-modal conflict in speech processing (*7-10*). Stimuli consisted of 40 high-frequency Chinese characters selected for familiarity, visual simplicity, and phonological clarity. Each stimulus was presented for 600 ms in both visual and auditory modalities, with pronunciations recorded from a standardized Mandarin phonetic database to ensure consistency. Trials were evenly divided into congruent (AVc) and incongruent (AVi) conditions (20 trials each) and presented in a randomized order.

The lexical decision task, by contrast, was designed to provide a window into the development of semantic recognition and evaluation, a core component of language comprehension. In this task, participants were presented with a pair of stimuli in each trial: a prime word followed by a target word. For each pair, they were instructed to judge the lexicality of each stimulus, indicating whether both the prime and the target were real words or pseudowords. The paradigm systematically manipulated the lexical status of both the prime and the target, yielding seven conditions: semantically related real–real, semantically unrelated real–real, associatively related real–real, associatively unrelated real–real, pseudoword–pseudoword, real–pseudoword, and pseudoword–real. A previous study using this paradigm has validated its behavioral efficacy, demonstrating robust effects of lexicality across all conditions (Ju Ping et al., submitted). For the purpose of the current investigation, which focuses on the developmental trajectories of semantic processing, we analyzed data pooled across two task conditions: real word-pseudoword condition and pseudoword-real word condition. Rather than comparing these two conditions to one another, we collapsed across them to directly contrast the processing of individual real words against pseudowords independent of their position within the stimulus pair. This fundamental item-level contrast has been shown to index increased lexical access demands, as pseudowords typically require additional verification processes during word recognition (*11, 12*). Stimuli consisted of high-frequency two-character Chinese words (e.g., “coffee”, “crow”, and “nose”) and pseudowords created by recombining components of real words into semantically implausible forms. Real and pseudoword stimuli were non-overlapping, with 44 trials per condition.

Together, these tasks tap into complementary levels of language processing, from perceptual integration to semantic evaluation, allowing us to trace the development of the language ability across different cognitive demands.

#### *1.3 Procedures*

The experiment was conducted in a quiet, well-lit room to ensure participant concentration. Stimuli were presented using E-Prime 2.0 software, and high-quality sound-isolating headphones were used to ensure clear auditory playback. Before the formal experiment began, the researcher explained the purpose, procedure, and potential risks of the study in detail to each participant. For minors, both the participant and their legal guardians were informed, and written informed consent was obtained.

In the audiovisual integration task, prior to the main task, each participant completed a practice session consisting of 8 trials to familiarize themselves with the procedure and task requirements. In each trial, a fixation point appeared in the center of the screen for 800 ms to prompt attention. Then, a Chinese character appeared in the center of the screen for 600 ms, and simultaneously, its pronunciation was played through the headphones for the same duration (600 ms). Participants were asked to judge whether the visual character and the auditory sound matched. If they matched, the participant pressed the "F" key; if not, the "J" key. The experimental procedure is illustrated in Fig. S5D.

In the lexical decision task, prior to the main task, each participant completed 12 practice trials to familiarize themselves with the task procedures. Each trial began with a fixation point displayed at the center of the screen for 300 ms to prompt attention. The fixation remained on screen as a two-character spoken word was presented via headphones. Following the auditory stimulus, a blank screen appeared for 500 ms, followed by a response cue. Participants were instructed to make a lexical decision based on the auditory input, pressing the "F" key for pseudowords and the "J" key for real words. The experimental procedure is illustrated in Fig. S5E.

***1.4 Data acquisition***

For the child and adolescent sample, structural and functional data were acquired using a GE Discovery MR750 3T scanner. During the scan, participants were instructed to fixate on a central point, remain still, refrain from systematic thinking, and stay awake. rs-fMRI data were collected for 6 minutes. The imaging parameters were: TR = 2000 ms, TE = 30 ms, flip angle = 90°, 33 axial slices acquired in an interleaved order, matrix size = 64 × 64, field of view (FOV) = 220 mm, voxel size = 3.5 × 3.5 × 4.2 $mm^3$, and 180 time points were obtained. The T1-weighted structural images were acquired with the following parameters: TR = 2600 ms, flip angle = 12°, matrix size = 256 × 256, and voxel size = 1 × 1 × 1 $mm^3$. For the adult sample, structural and functional data were collected using two different 3T MRI scanners: the Siemens Magnetom Prisma and the Siemens Magnetom TrioTim syngo MR B17.

In the audiovisual integration task, due to the inclusion of adult participants from different acquisition batches, two sets of scanning parameters were used. (1) For 58 participants scanned using a SIEMENS MAGNETOM TrioTim syngo MR B17 system, the functional imaging parameters were: TR = 2000 ms, TE = 30 ms, flip angle = 90°, 240 time points (8 minutes), 33 slices with a thickness of 3.8 mm, FOV = 220 mm, voxel size = 3.4 × 3.4 × 3.8 $mm^3$. T1-weighted anatomical images were acquired with: TR = 2530 ms, flip angle = 35°, matrix = 256 × 256, voxel size = 1 × 1 × 1 $mm^3$. (2) For the remaining 18 participants scanned using a SIEMENS MAGNETOM Prisma system, the functional imaging parameters were: TR = 2000 ms, TE = 30 ms, flip angle = 90°, 240 time points (8 minutes), 58 slices with a thickness of 2.5 mm, FOV = 210 mm, voxel size = 2.5 × 2.5 × 2.5 $mm^3$. T1-weighted anatomical images were acquired with: TR = 2300 ms, flip angle = 7°, matrix = 256 × 256, voxel size = 0.8 × 0.8 × 0.8 $mm^3$.

In the lexical decision task, fMRI data were collected using the same scanning parameters as in the audiovisual integration task. Among the adult participants, 52 were scanned using a SIEMENS MAGNETOM TrioTim syngo MR B17 system, while the remaining 18 were scanned using a SIEMENS MAGNETOM Prisma system.

#### *1.5 Data preprocessing*

The rs-fMRI data preprocessing was conducted using the DPARSF toolbox (*13*) (http://rfmri.org/DPARSF). The first 10 time points were discarded to ensure signal stabilization. Then, using the middle slice of the scan as the reference slice, slice timing correction was performed separately for the three datasets. After that, each of the three datasets was processed with the following steps. Head motion correction was applied using six motion parameters. Each participant's T1-weighted image was co-registered to the functional image, segmented into gray matter, white matter, and cerebrospinal fluid (*14*), and normalized to MNI space (*15*). FC analyses were band-pass filtered at 0.01–0.1 Hz. Participants with mean FD > 0.2mm were excluded; 6 participants were removed based on this criterion. Mean FD was included as a covariate in statistical analyses to control for head motion artifacts (*16*).

T1-weighted images were processed using FreeSurfer 7.4.1 (http://surfer.nmr.mgh.harvard.edu/) for cortical surface reconstruction and volumetric segmentation. The automated pipeline included skull stripping, gray matter segmentation, cortical surface reconstruction, surface inflation, spherical registration, parcellation, and surface-based metric extraction (*17-19*). Cortical thickness and surface area were computed at each vertex and then smoothed with a 10 mm full-width at half-maximum (FWHM) Gaussian kernel.

#### *1.6 Data analysis*

##### *1.6.1 Behavioural data analysis*

Participants with accuracy rates below 70% and reaction times (RTs) beyond three SDs from a participant's mean were excluded from further analysis (*20-22*). To model the developmental trajectory of RTs across age, a Generalized Additive Mixed Model (GAMM) was employed using the *mgcv* package in R (*23, 24*). Additionally, for each task, linear mixed-effects models (LMM) of RTs across two experimental conditions and three age groups were conducted in R to examine group differences (*25, 26*). Group and condition were included as fixed effects, with participants and items as random effects. Random slopes that failed to converge or resulted in overfitting were sequentially removed from the full model until convergence was achieved (*27, 28*).

In the audiovisual integration task, for each participant, mean RTs were computed separately for the AVc and AVi conditions. The RT difference was defined as the average RT in the incongruent condition minus that in the congruent condition, serving as a behavioral index of audiovisual integration ability (*29, 30*). In the lexical decision task, RTs were similarly calculated for the real-word and pseudoword conditions. The RT difference was defined as the mean RT for pseudowords minus that for real words, serving as an index of lexical decision efficiency (*11, 12*). To model the developmental trajectory of RT difference across age, a GAMM was also employed.

To further investigate which condition primarily contributed to the RT difference, correlation analyses were conducted between the RT difference and the mean RTs in each condition (congruent vs. incongruent in the audiovisual integration task; real-word vs. pseudoword in the lexical decision task). In addition, the RT difference was correlated with FC strength and structural measures (i.e., cortical thickness and surface area) to examine brain–behavior relationships.

*1.6.2 Brain network analysis*

We focused on two meta-analytically defined networks relevant to language, linked to audiovisual integration (AVI) and lexical decision (LD). For the audiovisual integration regions, ROIs were identified using NeuroSynth by searching the term "audiovisual". Consistent with previous findings (*30-33*), the resulting network primarily included the superior temporal regions bilaterally (see Fig. S6A). These functional regions were mapped onto the Glasser360 multimodal parcellation template (*34*) using a threshold of $Z > 4.5$ and voxel overlap >10%, yielding 7 ROIs (Fig. S6C). For the lexical decision network, ROIs were identified using the same procedure, based on a NeuroSynth meta-analysis for the term "lexical decision" (Fig. S6B). The resulting activation patterns were also overlaid onto the Glasser360 template using identical criteria ($Z > 4.5$, voxel overlap > 10%), resulting in 12 ROIs across left inferior frontal gyrus, supplementary motor area, temporal and parietal regions (Fig. S6D).

ROI-to-ROI analyses were conducted within the two networks, and each ROI was also used as a seed region for ROI-to-whole-brain connectivity analyses, with the AAL116 atlas serving as the template. FC analyses were implemented using the DPASF toolbox (http://www.restfmri.net). For each participant, the average time series within each ROI was extracted, and both ROI-to-ROI and ROI-to-AAL116 correlation matrices were computed. The resulting individual-level correlation maps ($r$-maps) were then Fisher $z$-transformed to obtain standardized $z$-FC maps, which were subjected to LMM analyses to examine developmental differences in FC. In addition, FC strength was correlated with cortical thickness and surface area of each ROI to further explore whether structural properties modulate the developmental changes in FC. All analyses were corrected for multiple comparisons using the Bonferroni method.

*1.6.3 Brain network–behavior analysis*

For both tasks, we extracted the mean timeseries for each participant from the ROIs within the audiovisual integration network and the lexical decision network, respectively, and computed both ROI-to-ROI and ROI-to-voxel correlation matrices. To control for false positives, multiple comparison correction for both ROI-to-ROI and ROI-to-voxel FC analyses was performed using Gaussian Random Field (GRF) theory (voxel-level $p < 0.005$, cluster-level $p < 0.05$). Significant FC strengths were further correlated with behavioral performance (i.e., RT differences). To account for potential site-related variability due to data acquisition across different scanners and institutions, scanning site was modeled using three binary dummy variables and included as covariates in all analyses. Additional covariates included mean framewise displacement (FD) as an index of head motion, gender, and age.

To examine the network distribution of FC results, we hypothesized that the regions sensitive to both the audiovisual integration and lexical decision tasks would show significant overlap with dorsal attention, ventral attention, and frontoparietal networks, which are recruited to support a wide variety of tasks as part of the multiple-demand system (*35-39*). Significant clusters from both tasks were overlaid onto the seven large-scale cortical networks defined by Yeo et al. (2011), using the Chinese HCP-derived 7-network parcellation(*40*), to assess the distribution of activation across networks.

*1.6.4 Individual deviation and dip contribution analyses*

To clarify the functional significance of the adolescent connectivity dip observed at the group level, we conducted two complementary analyses. First, we examined whether individual differences in the magnitude of the dip were behaviorally meaningful. For each age group, each participant's FC strength was expressed as a deviation (in standard deviation units) from the group mean. These deviation scores were then correlated with RT differences in each language task, allowing us to test whether individuals with a more pronounced dip (lower-than-average FC) or a weaker dip (higher-than-average FC) showed corresponding differences in performance.

Second, we assessed the system-level consequences of the dip by asking whether connections that weaken during adolescence still support overall network integration. To do this, connections were classified as "dip" connections if their FC strength was significantly lower in adolescence than in both childhood and adulthood; all remaining connections were classified as "non-dip". We then evaluated how each class of connections related to global network efficiency during adolescence, testing whether dip connections, despite their transient weakening, continued to contribute to integrative network organization. All analyses included scanning site, head motion (mean FD), gender, and age as covariates.

*1.6.5 Brain structure–behavior analysis*

To assess whether functional reorganization in adolescence was paralleled by a progressive uncoupling of behavior from structural metrics, we examined correlations between cortical thickness, surface area and RT difference. This analysis controlled the effects of site, mean FD, gender, and age as covariates. Furthermore, to control the family-wise error rate associated with multiple structural–behavioral tests, Bonferroni correction was applied separately within each task. Specifically, in the audiovisual integration task, 14 comparisons were corrected (7 regions × 2 structural metrics), whereas in the lexical decision task, 24 comparisons were corrected (12 regions × 2 structural metrics).

### 2 Supplementary Text

#### *2.1 Results of Two Behavioral Task*

In the audiovisual integration task, the GAMM results showed that RTs exhibited a significant age-related developmental trajectory in both the AVc (Fig. S3A) and AVi (Fig. S3B) conditions (AVc: $edf = 3.12$, $F = 3.37$, $p < .001$; AVi: $edf = 3.26$, $F = 3.48$, $p < .001$). An LMM of RTs under the AVc and AVi conditions across the three age groups is presented in Fig. S3C. The model revealed significant main effects of both condition and group. RTs were slower in the AVi than in the AVc condition (964 ± 365 ms vs. 897 ± 364 ms, $\beta = 94.13$, SE = 11.73, $t = 8.03$, $p < .001$), and adults responded faster (710 ± 266 ms) than adolescents (925 ± 365 ms, $\beta = -160.71$, SE = 23.07, $t = -6.97$, $p < .001$) and children (1157 ± 450 ms, $\beta = -395.61$, SE = 32.10, $t = -12.33$, $p < .001$). A significant interaction between condition and group [$F(2, 9750.6) = 5.68$, $p = .003$] indicated that the incongruence effect diminished with age: robust in children (1203 ± 450 ms vs. 1110 ms ± 451 ms, SE = 11.70, $z = -8.03$, $p < .001$) and adolescents (959 ± 365 ms vs. 891 ms ± 364 ms, SE = 9.71, $z = -7.08$, $p < .001$), but smaller in adults (730 ± 255 ms vs. 691 ± 276 ms, SE = 11.30, $z = -3.47$, $p < .001$). The GAMM further showed that the RT difference was significantly correlated with age (see Fig. S3D), showing a linear developmental trend ($edf = 1.00$,

$F = 4.32, p < .05$).

In the lexical decision task, the GAMM results showed that RTs also exhibited significant age-related developmental trajectories in both the real word (Fig. S3E) and pseudoword (Fig. S3F) conditions (real word: $edf = 2.83, F = 3.05, p < .001$; pseudoword: $edf = 2.66, F = 2.94, p < .001$). The LMM (Fig. S3G) revealed significant main effects of both condition and group. RTs were longer for pseudowords than real words (1230 ± 492 ms vs. 1122 ± 413 ms, $\beta = 90.83$, SE = 10.32, $t = 8.80, p < .001$), and adults again outperformed both adolescents (1194 ± 424 ms, $\beta = -112.43$, SE = 21.60, $t = -5.21, p < .001$) and children (1254 ± 594 ms, $\beta = -217.61$, SE = 32.95, $t = -6.61, p < .001$). The interaction between condition and group was marginal [$F(2, 21099.1) = 2.88$, $p = .056$], but simple effects confirmed a consistent pseudoword cost across age groups, found in children (1299 ± 626ms vs. 1208 ± 554 ms, SE = 10.30, $z = -8.80, p < .001$), adolescents (1248 ± 461 ms vs. 1140 ± 381 ms, SE = 8.86, $z = -12.15, p < .001$), and adults (1144 ± 255 ms vs. 1018 ± 276 ms, SE = 10.50, $z = -12.06, p < .001$). Unlike the audiovisual integration task, however, the GAMM results (Fig. S3H) revealed no significant linear or nonlinear relationship between RT difference and age: $edf = 1.00, F = 0.15, p = .70$.

To examine how RT differences relate to condition-specific RTs across age groups, we computed correlations separately for the audiovisual integration and lexical decision tasks. In the audiovisual integration task, during childhood (Fig. S4A), RT difference was significantly correlated with RT in both the congruent condition ($r = -0.37, p = .001$) and the incongruent condition ($r = 0.27, p = .021$). However, during adolescence (Fig. S4B), RT difference was significantly correlated only with RT in the incongruent condition (incongruent condition: $r = 0.34, p < .001$; congruent condition: $r = -0.18, p = .772$), and the same pattern was observed in adulthood (Fig. S4C; incongruent condition: $r = 0.68, p < .001$; congruent condition: $r = 0.03, p = .072$). A parallel pattern was observed in the lexical decision task. In childhood (Fig. S4D), RT difference was significantly correlated with RT in both the real-word condition ($r = -0.31, p = .017$) and the pseudoword condition ($r = 0.49, p < .001$). However, in adolescence (Fig. S4E) and adulthood (Fig. S4F), RT difference was significantly correlated only with RT in the pseudoword condition (adolescents: $r = 0.61, p < .001$; adults: $r = 0.73, p < .001$), with no significant correlations observed for the real-word condition (adolescents: $r = 0.07, p = .495$; adults: $r = 0.07, p = .550$). This convergence across tasks indicates that with age, performance differences are increasingly driven by processing demands in the more challenging condition, whereas easier conditions contribute less to the variability between conditions.

Although the condition × group interaction was not significant for either task (Fig. S3C and S3G), these correlation patterns suggest that in childhood, performance differences were associated with both the easier and the more demanding conditions, suggesting a lack of differentiation in processing demands. In adolescence and adulthood, however, performance differences were associated only with the more demanding condition (incongruent trials in the audiovisual integration task and pseudoword trials in the lexical decision task), indicating the emergence of a selective sensitivity to task difficulty, possibly reflecting the maturation of cognitive control and lexical processing mechanisms. However, as these findings are based on correlational analyses, further research is needed for validation.

#### *2.2 Detail results for large-scale brain networks overlap*

In the audiovisual integration task (Fig. 4B), the child group showed significant regions predominantly in the default mode network (DMN, 42%) and the auditory network (49%), with a smaller proportion in the attention networks (i.e., ventral attention network [VAN] and dorsal attention network [DAN], 9%). This suggests that audiovisual integration in children relies more on internal cognition and basic sensorimotor processes, with relatively limited engagement of flexible control systems. In the adolescent group, FC was more broadly distributed, involving the DMN (40%), control and attention networks (i.e., frontoparietal network [FPN], VAN, and DAN, 13%) and SMN (7%), along with notable engagement of the visual network (4%). This broader distribution reflects the recruitment of more complex and flexible control systems to meet the increasing cognitive demands of the task. The adult group demonstrated a high reliance on the control and attention networks (15%), suggesting that audiovisual integration becomes more automated and efficiently controlled by domain-general executive and attentional systems. Across all age groups, the involvement of the VAN increased progressively, from 1% in the child group to 8% in the adolescent group, and 11% in the adult group, highlighting the growing importance of attentional control in audiovisual integration processing.

Additionally, voxel counts were highest in the adolescent group (Fig. 4B), suggesting heightened neural activity during this period. Specifically, the chi-square test revealed a significant association between group and brain network, $\chi^2(12) = 3215.73$, $p < .001$. Further standardized residual analyses indicated that the adolescent group showed significantly higher than expected counts in several networks, particularly in the FPN ($z = 5.0$), the somatomotor network ($z = 12.5$), the VAN ($z = 3.4$), and the visual network ($z = 9.7$), whereas counts in the DMN ($z = –6.8$) and the DAN ($z = –12.6$) were lower than expected. In contrast, both the adult and child groups generally exhibited negative or neutral residuals across networks. More importantly, at the overall level, the total count across the seven networks in the adolescent group ($n = 16186$) was significantly higher than in the adult ($n = 3868$) and child groups ($n = 3444$).

In the lexical decision task, the child group (Fig. 4C) showed significant engagement primarily within the DMN (63%) and the control and attention networks (36%). Within the latter, the FPN was dominant (21%), followed by VAN (15%). This indicates that children's lexical decision is more reliant on intrinsic cognitive resources, with attentional networks being less involved. In the adolescent group, the control and attention networks became the largest contributor (49%), followed by the DMN (30%), SMN (17%) and the visual network (4%). Within the control and attention networks, the DAN (11%), FPN (20%), and VAN (18%) all showed substantial involvement. This reflects a transitional stage characterized by broader recruitment of attentional and executive systems, supported by multi-sensory contributions. In the adult group, the control and attention networks (86%) were the most prominent, with overwhelming dominance of the DAN (37%) and minimal contributions from the VAN (3%). This indicates that lexical decision in adults has become highly specialized and goal-directed, relying primarily on the DAN. Overall, across age groups, the DAN showed a clear developmental increase, from 0% in children to 11% in adolescents, and reaching 46% in adults, highlighting its progressively central role in lexical decision-making.

Additionally, voxel counts were highest in the adolescent group (Fig. 4C), suggesting heightened neural activity and extensive recruitment of neural circuits during this developmental stage. The chi-square test revealed significant differences among the three groups, $\chi^2(12) = 5510.17$, $p < .001$. The adolescent group showed higher than expected counts in the DMN ($z =$

3.3), somatomotor network ($z = 9.6$), VAN ($z = 8.1$), and visual network ($z = 6.4$), whereas the adult group was primarily overrepresented in the DAN ($z = 47.3$).

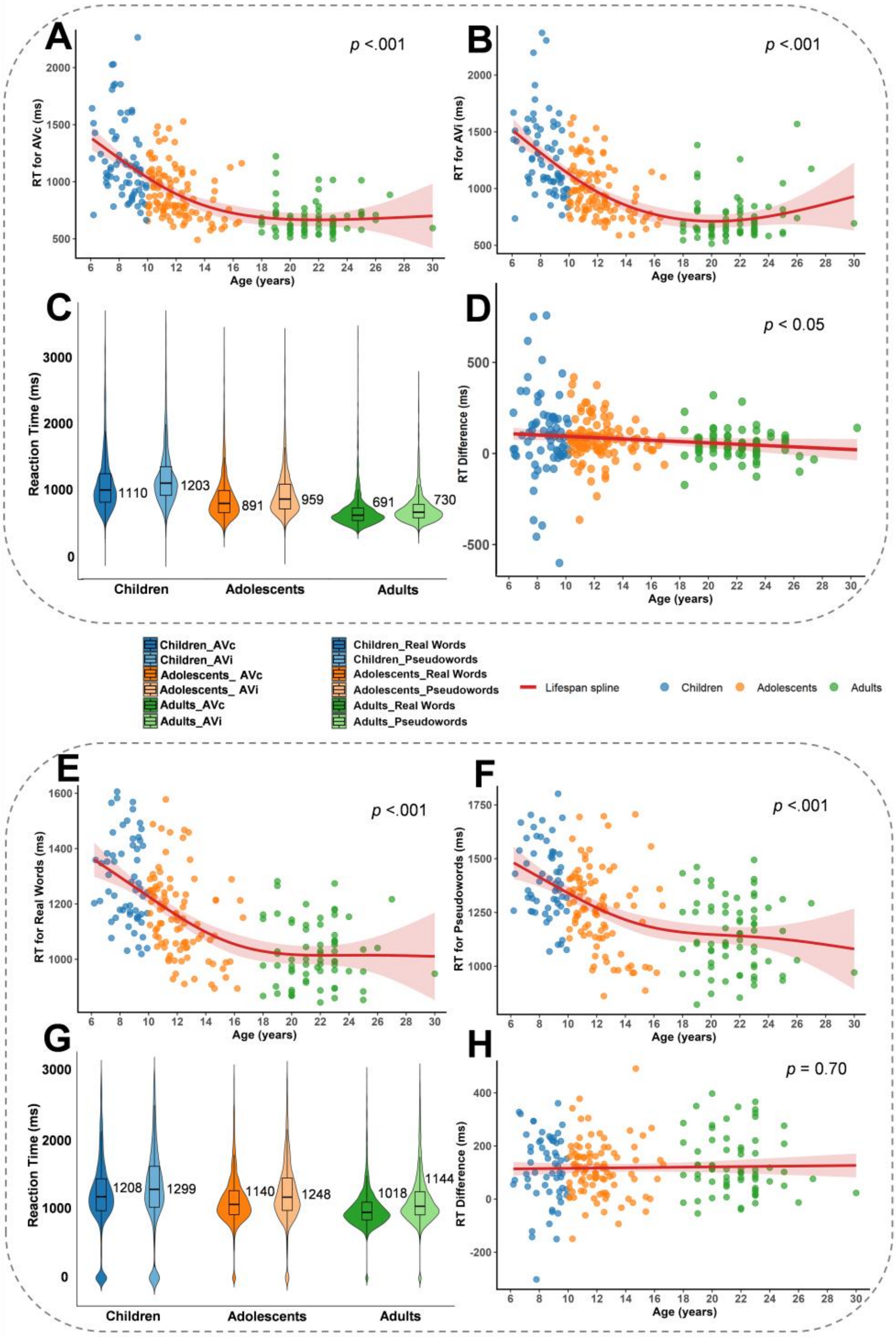


**Fig. S1. Behavioral results and developmental trajectories in the audiovisual integration and lexical decision tasks.** (**A**) and (**B**) show the age-related developmental trajectories of RTs in the AVc and AVi conditions, respectively. (**C**) LMM results of RTs across three age groups under congruent and incongruent audiovisual conditions. (**D**) Developmental trajectory of audiovisual integration across age groups. (**E**) and (**F**) show the age-related developmental trajectories of RTs in the real-word and pseudoword conditions, respectively. (**G**) LMM results of RTs across three age groups for real-word and pseudoword conditions in the lexical decision task. (**H**) Developmental trajectory of lexical decision performance across age groups. *Note*: Numbers in the violin plot indicate mean RTs. AVc = audiovisual congruent; AVi = audiovisual incongruent.

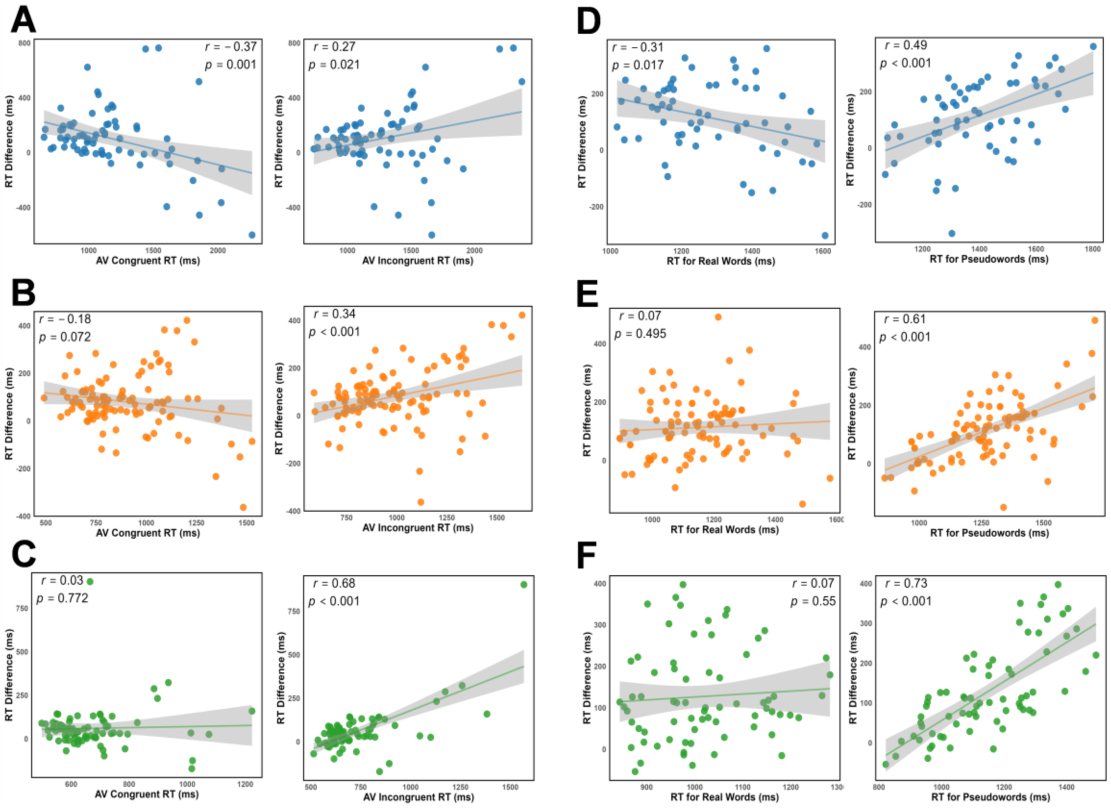


**Fig. S2. Correlations between the RT difference and condition-specific RT within each age group.** Panels (**A**), (**B**), and (**C**) show results for children, adolescents, and adults, respectively, in the audiovisual integration task. Panels (**D**), (**E**), and (**F**) show corresponding results for children, adolescents, and adults, respectively, in the lexical decision task.

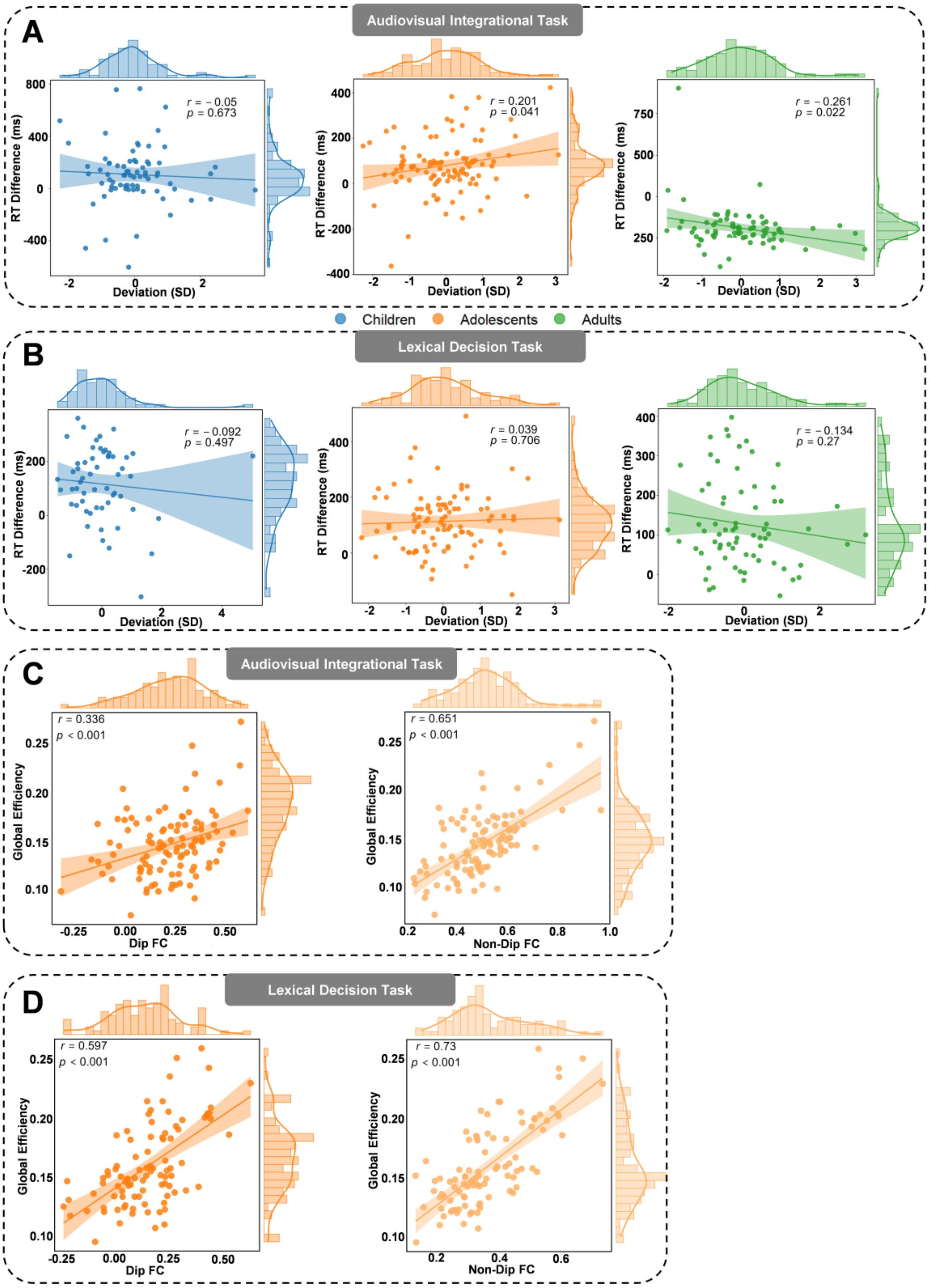


**Fig. S3. Relationships between individual deviation, behavioral performance, and adolescent dip FC.** Panel **A** and **B** show the correlations between individual deviation and behavioral performance for the audiovisual integration task and the lexical decision task, respectively. Panel **C** and **D** respectively show the correlations between dip and non-dip connections and global efficiency among adolescents during the audiovisual integration task and the lexical decision task.

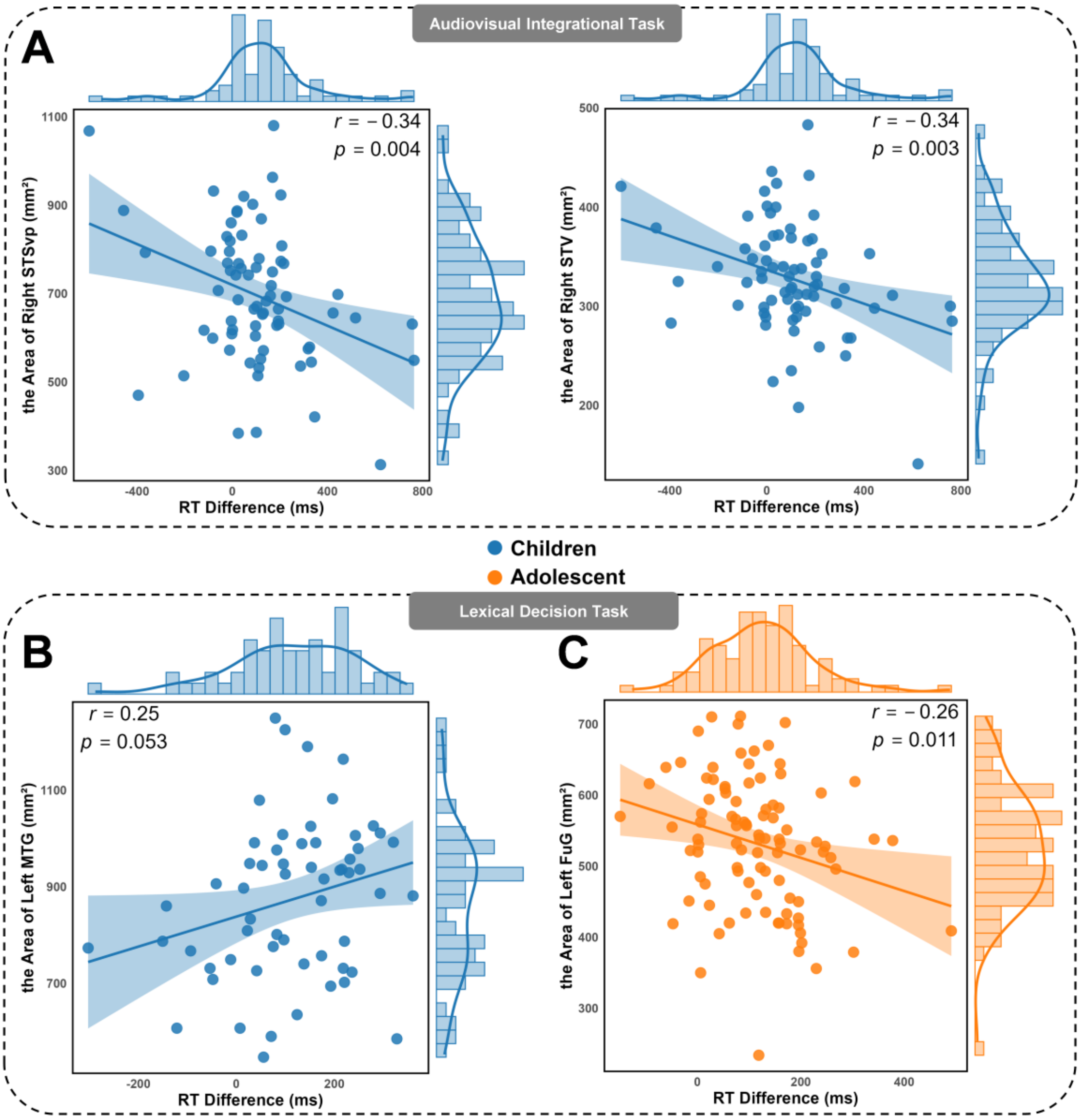


**Fig. S4. Correlations between structural metrics and behavioral performance.** Panel **A** shows the correlations between structural metrics and RT difference in childhood for audiovisual integration task. Panel **B** and **C** show the correlations between structural metrics and RT differences in the lexical decision task, presented separately for children and adolescence.

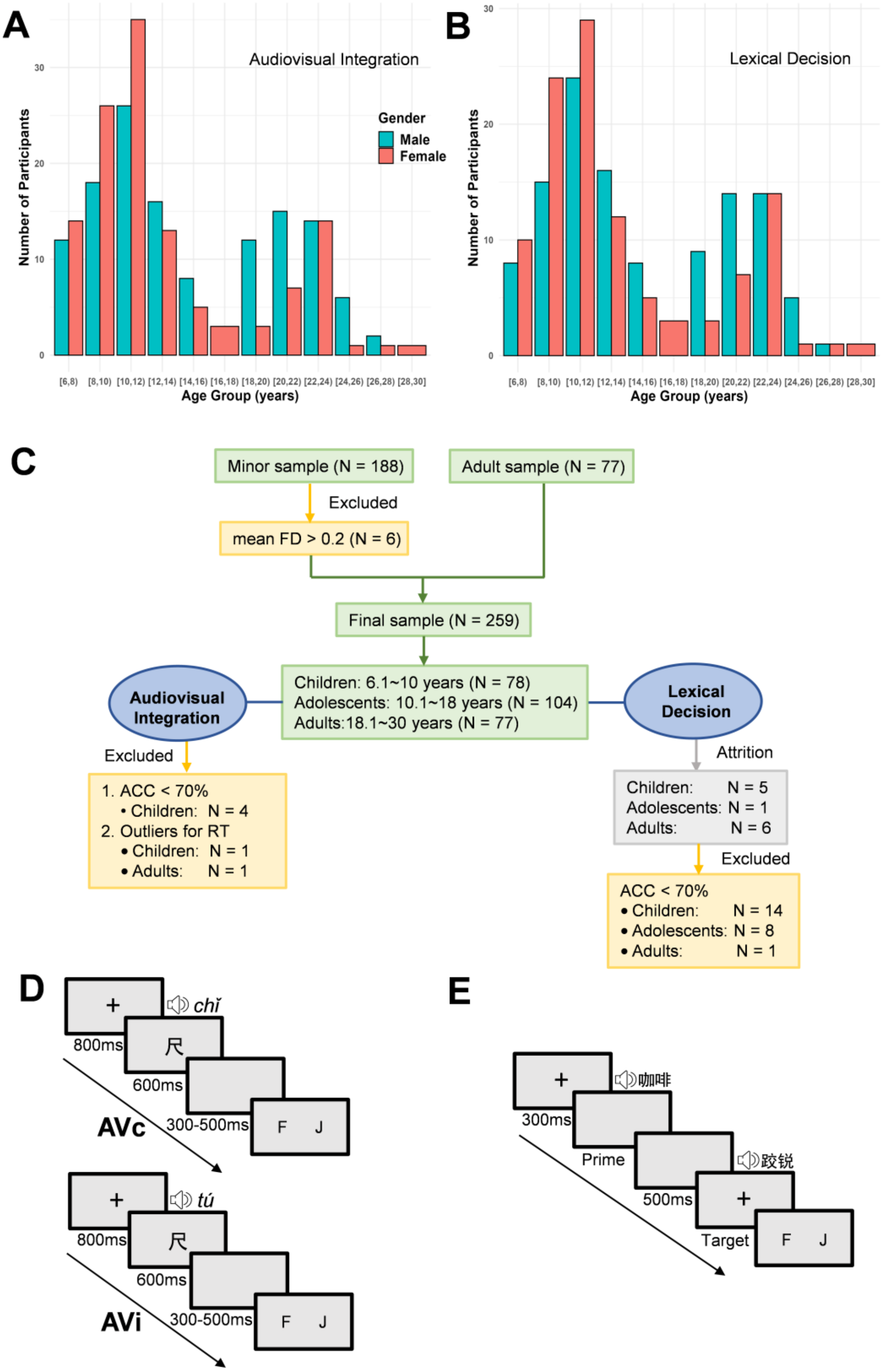


**Fig. S5. Overview of participants and and experimental procedures.** Age distribution of participants in the audiovisual integration task (**A**) and the lexical decision task (**B**). (**C**) Flowchart illustrating the participant inclusion process for each task. Experimental procedures for the audiovisual integration task (**D**) and the lexical decision task (**E**), respectively. *Note*: AVc = audiovisual congruent; AVi = audiovisual incongruent; RW = real word; PW = pseudoword.

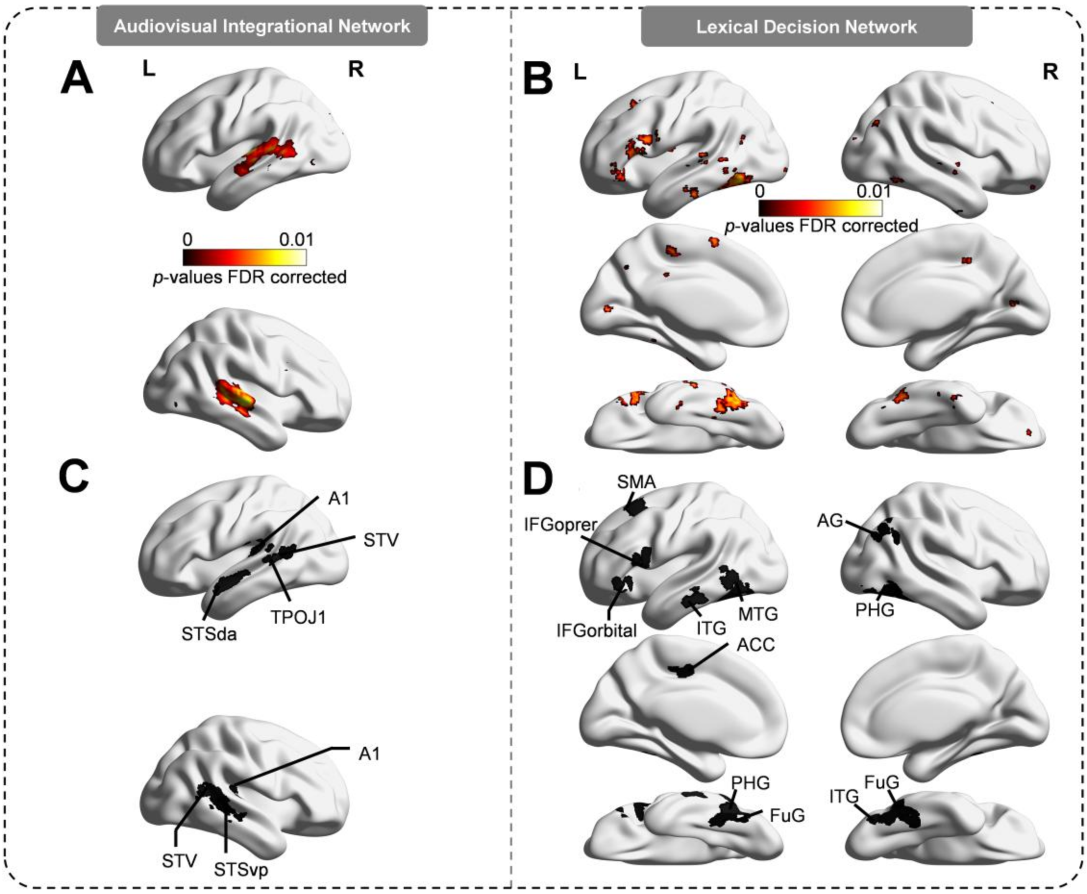


**Fig. S6. Meta-analytic and ROI-based neural networks for audiovisual integration and lexical decision.** Meta-analytic brain networks associated with the terms "audiovisual" (**A**) and "lexical decision" (**B**), identified using the NeuroSynth database. These maps reflect brain regions consistently activated across multiple studies using the respective search terms. (**C**) Seven core ROIs implicated in audiovisual integration, including the left primary auditory cortex (A1), superior temporal visual area (STV), temporo-parieto-occipital junction area 1 (TPOJ1), and dorsal anterior superior temporal sulcus (STSda), as well as the right A1, STV, and posterior ventral superior temporal sulcus (STSvp). (**D**) Twelve ROIs associated with the lexical decision task, comprising the left supplementary motor area (SMA), pars opercularis of the inferior frontal gyrus (IFGoprer), orbital part of the inferior frontal gyrus (IFGorbital), inferior temporal gyrus (ITG), middle temporal gyrus (MTG), anterior cingulate cortex (ACC), parahippocampal gyrus (PHG), and fusiform gyrus (FuG), as well as the right angular gyrus (AG), PHG, FuG, and ITG.

**Table S1. Summary of behavioral results in audiovisual integration and lexical decision tasks across age groups.**

| Task | Condition Effect | Age Group Effect | Condition × Age Interaction | Correlation Pattern | Age Trend |
|---|---|---|---|---|---|
| Audiovisual Integration | Incongruent > Congruent | Children > Adolescents > Adults | $p$ = .003 | Children: both conditions; Adolescents/Adults: incongruent condition only | Linear increase |
| Lexical Decision | Pseudoword > Real-word | Children > Adolescents > Adults | $p$ = .056 | Children: both conditions; Adolescents/Adults: pseudoword condition only | NS |

*Note*: "NS" indicates effects that were not statistically significant.

**Table S2. Overlap between significant brain regions and CHCP-derived Yeo seven large-scale networks.**

| | | Brain Regions | Visual | Somatomotor | Dorsal Attention | Ventral Attention | Auditory | Frontoparietal | Default Mode |
|---|---|---|---|---|---|---|---|---|---|
| Audiovisual Integration | Children | L.SPL | - | - | 38.43% | - | - | - | - |
| | | L.TPOJ1 | - | - | 0.11% | 0.82% | 44.39% | - | 38.24% |
| | Adolescents | L.CG | - | 52.32% | - | 40.79% | - | - | - |
| | | L.PostCG | - | 99.24% | 0.61% | - | - | - | - |
| | | L.MTG | - | - | 2.09% | - | - | 22.18% | 53.77% |
| | | L.MOG | 90.85% | - | - | - | - | 3.01% | 0.68% |
| | | L.STV | - | - | 0.22% | 11.92% | 39.92% | 0.37% | 43.08% |
| | | L.TPOJ1 | - | - | 0.11% | 0.82% | 44.39% | - | 38.24% |
| | | R.MTG | - | - | - | - | - | 62.50% | 27.02% |
| | | R.A1 | - | 7.64% | - | 0.37% | 88.16% | - | - |
| | | R.STV | - | - | - | 11.73% | 27.80% | 0.98% | 55.29% |
| | Adults | L.SFG | - | - | - | 0.69% | - | 11.58% | 31.88% |
| | | R.STV | - | - | - | 11.73% | 27.80% | 0.98% | 55.29% |
| Lexical Decision | Children | L.MFG | - | - | - | - | - | - | 62.96% |
| | | L.CG | - | 1.89% | - | 17.84% | - | 25.70% | 44.90% |
| | Adolescents | L.LG | 50.19% | - | - | - | - | - | - |
| | | L.IPL | - | 21.06% | 68.78% | - | - | 1.38% | - |
| | | L.IPL | - | - | 1.67% | 4.90% | - | 77.06% | 0.12% |
| | | L.PostCG | - | 87.96% | 0.81% | - | - | 2.43% | - |
| | | L.PostCG | - | 82.15% | 8.16% | - | - | 0.67% | - |
| | | L.PostCG | - | 65.19% | 27.30% | - | - | 1.16% | - |
| | | L.Insula | - | 0.20% | - | 40.90% | 0.12% | 27.62% | 1.25% |
| | | L.SMG | - | - | - | - | - | 3.05% | 87.36% |
| | | L.MFG | - | - | 0.30% | 3.70% | - | 26.07% | 69.19% |
| | | L.MFG | - | - | - | 0.18% | - | 2.67% | 68.81% |
| | | L.PreCG | 3.22% | - | 29.22% | - | - | 31.50% | 32.11% |
| | | R.IPL | - | - | 58.64% | - | - | 41.36% | - |
| | | R.MFG | - | - | 10.58% | - | - | 88.89% | - |
| | | R.PostCG | - | 64.15% | 15.84% | 2.84% | - | - | - |
| | | R.CG | - | 11.88% | 0.55% | 64.03% | - | 2.98% | 0.04% |
| | | R.Insula | - | 1.02% | - | 53.51% | - | 4.10% | - |
| | | R.MFG | - | - | - | 40.47% | - | 55.47% | - |
| | | R.Angular | - | - | - | - | - | 12.56% | 74.56% |
| | Adults | L.ITG | 0.93% | - | 41.01% | - | - | 2.51% | 36.90% |
| | | L.FuG | - | - | 77.44% | - | - | 22.39% | - |
| | | R.ITG | - | - | 23.77% | - | - | 76.23% | - |
| | | R.PostCG | - | 24.35% | 52.86% | 13.69% | - | - | - |
| | | R.IPL | - | - | 26.18% | - | - | 62.54% | - |

*Note*: “-” indicates no overlap.
